# Stabilized Independent Component Analysis outperforms other methods in finding reproducible signals in tumoral transcriptomes

**DOI:** 10.1101/318154

**Authors:** Laura Cantini, Ulykbek Kairov, Aurélien de Reyniès, Emmanuel Barillot, François Radvanyi, Andrei Zinovyev

## Abstract

**Motivation:** Matrix factorization methods are widely exploited in order to reduce dimensionality of transcriptomic datasets to the action of few hidden factors (metagenes). Applying such methods to similar independent datasets should yield reproducible inter-series outputs, though it was never demonstrated yet.

**Results:** We systematically test state-of-art methods of matrix factorization on several transcriptomic datasets of the same cancer type. Inspired by concepts of evolutionary bioinformatics, we design a new framework based on Reciprocally Best Hit (RBH) graphs in order to benchmark the method’s reproducibility. We show that a particular protocol of application of Independent Component Analysis (ICA), accompanied by a stabilisation procedure, leads to a significant increase in the inter-series output reproducibility. Moreover, we show that the signals detected through this method are systematically more interpretable than those of other state-of-art methods. We developed a user-friendly tool BIODICA for performing the Stabilized ICA-based RBH meta-analysis. We apply this methodology to the study of colorectal cancer (CRC) for which 14 independent publicly available transcriptomic datasets can be collected. The resulting RBH graph maps the landscape of interconnected factors that can be associated to biological processes or to technological artefacts. These factors can be used as clinical biomarkers or robust and tumor-type specific transcriptomic signatures of tumoral cells or tumoral microenvironment. Their intensities in different samples shed light on the mechanistic basis of CRC molecular subtyping.

**Availability:** The BIODICA tool is available from https://github.com/LabBandSB/BIODICA.

**Supplementary information:** Supplementary data are available at *Bioinformatics* online.

## 1 Introduction

Large-scale cancer genomics projects, such as The Cancer Genome Atlas (TCGA) and the International Cancer Genome Consortium (ICGC), are generating an overwhelming amount of omics data from multiple platforms and 10× Genomics released a scRNAseq dataset composed of 1.3 million mouse brain cells (Villani *et al.*, 2017). The available data offer us the unpredicted opportunity to understand cancer, its phenotypic properties, onset, progression and response to treatment. On the other hand, the massive sample size and high dimensionality of the current genomic data opens to all the computational and statistical challenges typical of “Big Data”. New computational and statistical paradigms are thus essential to leverage the full power of high-throughput data, including bulk transcriptomic, epigenetic, proteomic and single-cell data.

The use of matrix factorization (MF) approaches, reducing the dimension of the high-dimensional data into low dimensional subspaces, represents a powerful solution to this problem. The origins of MFs predate modern genomics. However, they are having a great success in biology because the state of a biological sample, such as a tumor sample, reflects multiple concurrent biological factors, from cell-type specific features to dynamic processes such as cell cycle. High-throughput data can be thus interpreted as a quantitative estimation of this mixture of biological factors together with technical noise due to samples processing and data generation. The deconvolution of these factors through matrix factorization approaches can thus provide an insightful interpretation of high-throughput data and increase our understanding of cancer. Also because each gene in MF has a chance to contribute to several hidden factors, unlike standard clustering methods.

The current state-of-art in the field of matrix factorization is represented by Principal Component Analysis (PCA), Non Negative Matrix Factorization (NMF) and Independent Component Analysis (ICA). Despite similar formulation of the approximation problem, each algorithm constrained in its own way and being applied to the same dataset, can lead to different sets of hidden signals (Zinovyev *et al.*, 2013). The existing differences among the various MFs impede to perform a one-to-one mapping between the factors identified by the different methodologies. A more sophisticated solution is thus required to compare state-of-art MFs in biology and, to our knowledge, no such framework has been developed till so far.

In this manuscript we developed a new metaanalysis method, based on exploiting Reciprocal Best Hit (RBH) relations between metagenes and clustering the RBH network. The method allows extracting robustly reproducible and biologically relevant features without imposing thresholds on the correlation coefficients unlike standardly used correlation networks. We also suggest a set of relevant criteria for benchmarking the matrix factorization methods used to produce the RBH network. We took advantage of multiple publicly available transcriptomic datasets from the same cancer type to test the reproducibility of the results obtained by different approaches. The framework that we here proposed and our use of publicly available data guarantees that any future methodology can be easily compared against those considered in this paper.

In general, we illustrated that there are marked differences between the various MFs. Our main result is in that stabilized ICA, a particular protocol of applying ICA to transcriptomic data, drastically outperformed alternative approaches. Using this method, and taking advantage of transcriptomic data available from tumor fragments, single-cells, Patient-Derived CRC Xenograft Models (PDX) and Liver Metastasis (LM), we mapped a landscape of independent hidden factors shaping colorectal cancer transcriptomes.

## 2 Methods

### 2.1 Description of the considered state-of-art MF approaches

The general idea behind Matrix factorization algorithms is to reduce data dimensionality through a decomposition approach. Given the natural representation of high-dimensional biological data as a matrix of measurements (expression counts, methylation levels, protein concentrations, etc) with different samples represented in the columns and different molecules of interest (genes, proteins, etc.) represented in the rows. MFs decompose such a matrix X (n × m) into the product of an unknown mixing matrix A (n × k) and an unknown matrix of source signals S (k × m). In the following, we denote the columns of the molecule-level matrix “metagenes” and the rows of the sample-level matrix “metasamples”. Metasamples and metagenes are learned based upon the assumption that the number *k* of biological factors occurring in the input dataset is smaller than either the number of rows or columns in the input matrix (X). Determining the optimal number *k* of biological factors to use in the factorization is critical to its interpretation. The appropriate selection depends upon the algorithm and is an active area of research (Kairov *et al.*, 2017).

We give here a brief summary of the MFs that we compare in the present work for detecting low-dimensional biological factors from large-scale transcriptomic data sets. For more detailed description the reader is referred to the original publications.

#### Principal Component Analysis (PCA)

PCA maximizes variance and it leads to orthogonal factors. Because PCA learns features that explain most of the variation in the data, it conflates multiple biological processes into single components and thus it is not the optimal approach to learn the specific genes co-regulated by a specific biological factor (Ochs and Fertig, 2012). In this regard, PCA may be of use in experimental paradigms in which the processes or conditions of interest represent the strongest sources of variation, such as to remove technical artifacts (Parker *et al.*, 2014; Martignetti *et al.*, 2016). The limitations of PCA in application to transcriptomic data was highlighted in many publications (Saidi *et al.*, 2004).

#### Non Negative Matrix Factorization (NMF)

NMF solves the minimum representation error under the constraint of having all the elements of the A and S matrix non-negative (Lee and Seung, 1999; Ochs *et al.*, 1999). Such constraint of the factors learned by NMF matches the non-negative nature of transcriptomic data (e.g., read counts) it is thus considered as a natural approximation. Moreover NMF allows existence of coupled factors thereby modeling coregulation (Moloshok *et al.*, 2002).

#### Independent Component Analysis (ICA)

ICA has been originally proposed in the context of signal processing to decompose a multivariate signal into factors characterized by non-Gaussian distributions that are as independent as possible (Hyvärinen *et al.*, 2001). Minimizing statistical dependence ensures that the patterns learned by ICA come from distinct biological processes. Different protocols to apply ICA exist and currently no standard approach exists. The main difference in the existing approaches concerns what is considered as source signal matrix in the decomposition. Indeed some aims at maximizing the non-gaussianity of metagenes (Biton *et al.*, 2014; Kairov *et al.*, 2017; Kong *et al.*, 2008; Lee and Batzoglou, 2003), while others maximize non-gaussianity of metasamples (Meng *et al.*, 2016; Barillot, 2013). We here compared both approaches. We will call in particular “Stabilized ICA” the protocol previously proposed by us that maximizes kurtosis of metagenes and searches for stable components by performing a bootstrap approach and thus is able to prioritize stable components (Biton *et al.*, 2014; Kairov *et al.*, 2017). We will instead denote with “ ICA’ ” the application of ICA that maximizes kurtosis of metasamples, which corresponds to apply ICA to the transposed expression matrix, operation that in MATLAB is denoted with the symbol “ ’ ”, from which the choice of the name is originated. For a typical transcriptomic dataset analysis, the advantages of Stabilized ICA are: a) having the number of objects larger than the number variables improves convergence of the algorithm and stability of the resulting components; b) resulting metasamples (unlike metagenes) might appear strongly correlated, reflecting coupling of certain biological mechanisms. The detailed description of the stabilized ICA protocol with exact definition of the stability measure is provided in (Kairov *et al.*, 2017).

To compare such approaches we need to define a computational framework and to choose a biological context on which perform the comparison. These aspects are discussed in the next sections.

### 2.2 Biological context and datasets chosen for the comparison

We here chose colorectal cancer (CRC) as biological context for our comparison. The choice of working in a cancer context is due to the fact that cancer is characterized by wider transcriptomic variations. In addition, a lot of transcriptomic datasets are already available and carefully annotated in cancer biology. Our framework takes advantage from having several independent transcriptomic datasets for the same cancer type. Among the various cancers, CRC is one of the most common cancer both in men and women and it is the one for which 14 independent and large transcriptomic data sets could be easily downloaded. Indeed, strong efforts have been done by multiple research groups to identify CRC subtypes, i.e. groups of CRC samples with homogeneous molecular and/or clinical characteristics.

Recently a consensus CRC subtyping (CMS) has been proposed (Guinney *et al.*, 2015). In this work a collection of all the transcriptomic datasets previously employed for subtyping has been made available to the public, this is the resource that we employed for our comparison. For details concerning the datasets see Supplementary Table 1.

Moreover, to test the effects of the platform on our analysis we also used 4 TCGA ovarian cancer datasets profiled with various microarrays technologies: Affymetrix U133, Agilent and Affymetrix HuEx, plus RNAseq. These 4 datasets have a total of 418 common samples. They are thus the perfect resource to test how the choice of the platform affects the different MFs.

### 2.3 Computational framework for metagene comparison

We defined a new framework to compare four state-of-art MF algorithms: PCA, NMF, ICA’ and stabilized ICA (see Figure 1 for a schematic representation of the framework). The challenge that we had to face here was to standardize the different methods in order to make them comparable. First, the number *k* of components in which the expression matrix is decomposed should be chosen for all the compared MFs. We overdecomposed the matrices and we fixed the same number of components for all the MFs (see Supplementary Table 1 for details). Overdecomposition here stands for the fact that the selected number of components is taken much larger than the estimation of the effective transcriptome dimension estimated used one of the existing approaches (Keiser rule, broken-stick distribution rule, Maximally Stable Transcriptomic Dimension approach).

**Fig. 1.**
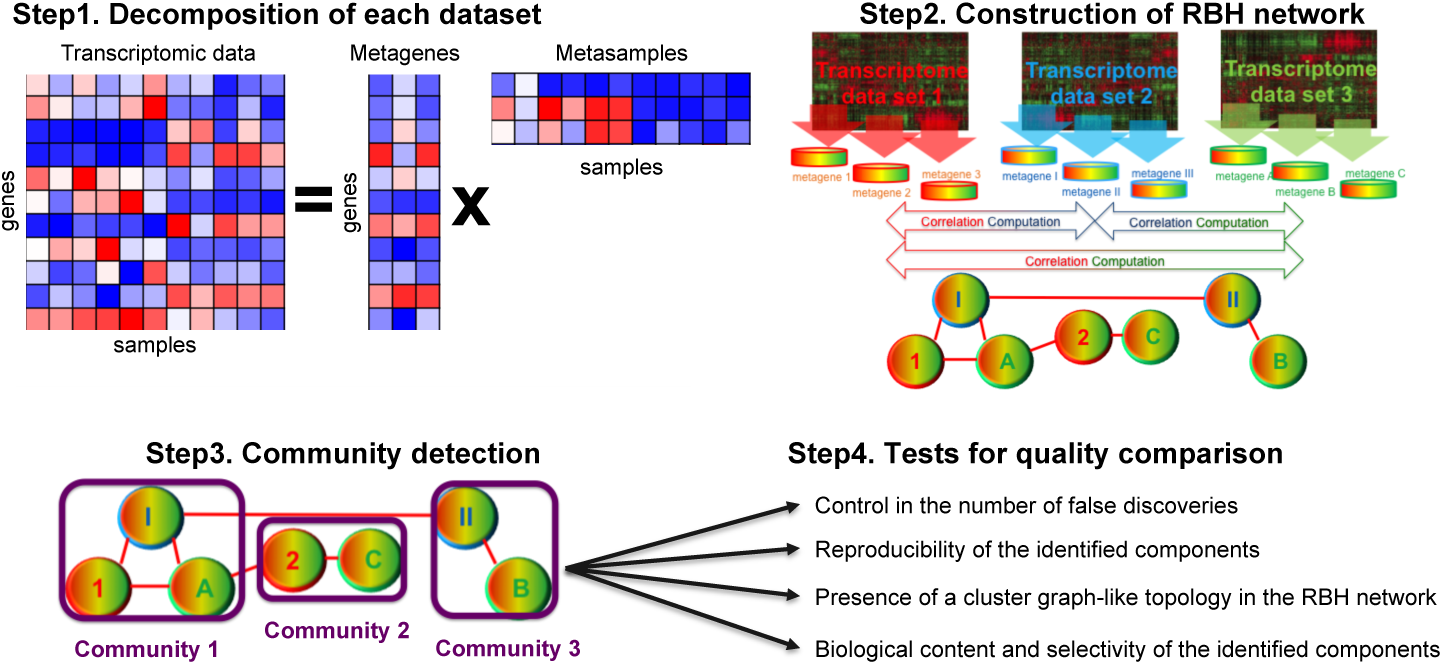
Schematic representation of MF comparison framework.

In our previous work, we have shown that in case of ICA, overdecomposition is not detrimental for the interpretability of the resulting components (Kairov *et al.*, 2017). The same is evidently true for PCA, since the resulting components are orthogonal. For NMF the number *k* of components in which a dataset should be decomposed is frequently decided by looking at the last local maximum of the cophenetic coefficient, summarizing the results of a consensus over different runs of the algorithm (Brunet *et al.*, 2004). We thus chose to also compare our four algo-rithms against the version of NMF whose number of components is chosen based on the cophenetic coefficient, called in the following “cophNMF”. Such comparison is reported in Supplementary Table 2.

As shown in Figure 1 our new framework is composed of 4 main steps to be separately performed for each MF algorithm. The only inputs required to perform the comparison are as many independent transcriptomic datasets as possible for the same biological context. In our case, these were the 14 CRC transcriptomic datasets from (Guinney *et al.*, 2015).

At step 1, each dataset is decomposed into a set of metagenes and metasamples. One of the main methodological novelties of this work is represented by step 2, where the graph of univocal correspondences between the metagenes obtained from the various independent datasets is reconstructed. Considering two transcriptomic datasets (T^1^ and T^2^), we define as best-hit of a metagene (M^1^_i_) of dataset T^1^ the metagene (M^2^_n_) of datasets T^2^ if the absolute correlation between M^1^_i_ and M^2^_n_ is maximal in respect to the correlations of M^1^_i_ with all the metagenes of dataset T^2^. Therefore, to have a link in our graph between M^1^_i_ and M^2^_n_ we require such condition to be verified in both directions, meaning that M^2^_n_ has to be the best-hit for M^2^_n_ among all the metagenes of dataset T^2^ and M^1^_i_ has to be the best-hit for M^2^_n_ among all the metagenes of dataset T^1^. Here and in the following we will refer to the obtained graph as Reciprocal Best Hit (RBH) graph. This name is chosen in analogy with the namesake most common working definition of orthology in comparative genomics. Essentially, when the proteins encoded by two genes, each in a different genome, find each other as the best scoring match in the other genome the two genes are considered to be orthologous according to the RBH procedure (Tatusov *et al.*, 1997; Bork *et al.*, 1998). The idea behind our approach is thus to identify orthologous biological factors across different transcriptomic datasets. This approach is conceptually very different from the frequently used approach of constructing a correlation graph. Indeed when using correlation a complete graph is reconstructed. To extract information from it some links need to be filtered out. Different filtering approaches exist, going from a simple thresholding to more sophisticated strategies (Serrano *et al.*, 2009). Moreover, once the optimal filtering algorithm has been chosen, also the value of the threshold needs to be selected. The obtained results are thus dependent on both the chosen filtering approach and threshold value. On the opposite, RBH is free of threshold choice and it leads to relatively sparse graphs. In Supplementary Figure 1 we compare the number of RBHs and the dimension of the largest connected component of the correlation network for various thresholds vs. the RBH network in all the MFs.

Following the reconstruction of the RBH graph, we observed that the components detected by NMF were strongly biased toward the genes’ average expression (see Supplementary Figure 2), i.e. the vector containing in each row the average expression of a gene across all the samples of the dataset. This resulted in a RBH graph where most of the metagenes are strongly correlated with each other, which makes the comparison with other MFs impossible. As a further standardization, we thus corrected the metagenes obtained with all the MFs for the genes’ average expression before constructing the RBH graph. We performed this task by regressing the metagenes over the genes’ average expression and using the residues of such regression to compute the RBHs.

At step 3, RBH graph communities are detected using Markov Clustering algorithm (MCL), this is another novelty in respect to previous works (Biton *et al.*, 2014; Kairov *et al.*, 2017). The communities reflect existence of biological factors strongly reproduced across different CRC transcriptomes. Finally, at step 4 different objective measures are computed to compare the results obtained by the various MFs. The idea in this last step is to evaluate the performances of the different algorithms focusing on measures that are of practical interest to researchers when analyzing high-throughput data. In particular, we evaluated the ability of the different MFs to (i) control the number of false discoveries; (ii) determine reproducible components; (iii) derive an RBH graph characterized by tight community structure and (vi) identify biologically meaningful and selective components, meaning components able to accurately and univocally predict known biological signals. The results of these tests are extensively discussed in the results section.

### 2.4 Interpretation of the communities obtained in the RBH graph

Once defined the best performing MF algorithm, we investigated which already known and new insights could be obtained using our newly proposed framework. With this aim we added to the analysis other four datasets: single-cell RNAseq from normal and tumoral CRC tissue (Li *et al.*, 2017), Patient-derived Xenograft (PDX) CRC Models transcriptomic data and liver metastasis (LM) transcriptomic data (Isella *et al.*, 2017). Given the heterogeneity of such data in respect to the previous 14 we only employed them for the biological characterization and not in the assessment of MF algorithm performances. We characterized the communities obtained in the RBH graph with all the available biological annotations: MsigDB signatures(Liberzon *et al.*, 2011), cell types signatures (Aran *et al.*, 2017), tissue-specific Transcription Factor (TF)-target associations (Marbach *et al.*, 2016), ToppGene (Chen *et al.*, 2009), the clinical annotations available for the various transcriptomic datasets, the CMS and CRIS subtypes (Guinney *et al.*, 2015; Isella *et al.*, 2017) and finally the cell types associations available from the single-cell RNAseq data (Li *et al.*, 2017).

We employed the metasamples of all the components contained in a community to test the association with clinical, CMS and CRIS annotations. We tested the significance of such annotation by performing a two-sided Wilcoxon or Kruskal-Wallis depending if the comparison was involving two classes (such as gender) or more than two (such as the 4 CMS subtypes), respectively. If a community contained a single-cell derived component we tested its association with a specific cell type with a KruskalWallis test. We considered as significant those tests having a Benjamini-Hochberg corrected p-value lower than 0.05.

For all the other biological annotations involving genes we employed the metagenes contained in each community. We thus associated to each community of the RBH graph a “meta-metagene” corresponding to the average of all the metagenes contained in the community, paying attention to first concordantly orientate all the metagenes of the community based on the signs of their correlations. We then used Preranked GSEA with MsigDB signatures to test the association of our metagenes with specific pathways and biological functions (Subramanian *et al.*, 2005). We then defined as top-contributing genes of a community those genes having a weight in the meta-metagene higher than 3 standard deviations in absolute value. We tested for the intersection of the top-contributing genes with cell types specific signatures using a Fisher’s exact test and we applied ToppGene to them. Finally, to detect possible TFs regulating the communities we used the tissue-specific TF-target associations in (Marbach *et al.*, 2016) and tested for the presence of a TF and a significant number of its targets (according to a Fisher’s exact test) in the top-contributing genes of each meta-metagene.

### 2.5 BIODICA tool for computing and interpreting the stabilized independent components

We have developed and released to public BIODICA (ICA applied to BIOlogical Data) tool, available from https://github.com/LabBandSB/BIODICA. BIODICA implements the protocol of Stabilised ICA to transcriptomic and other omics data. BIODICA provides both a command line and a user-friendly Graphical User Interface (GUI) for high-performance ICA analysis, including bootstraping and further stability analysis. BIODICA is based on previous high-performing implementations of fastICA and icasso algorithms, allowing to boost the ICA computation time by an order of magnitude compared to the existing fastICA implementation in R. BIODICA provides the possibility to run ICA decomposition of several orders and it computes the Maximally Stable Transcriptome Dimensionality (MSTD) measure, which can be used to determining the optimal number of independent components (Kairov *et al.*, 2017).

Moreover, BIODICA provides several tools for downstream interpretation of the resulting metagenes and metasamples. This interpretation includes standard functional analyses such as Preranked Gene Set Enrichement Analysis (Subramanian *et al.*, 2005) and hypergeometical tests using automated ToppGene web-service (Chen *et al.*, 2009), but also a built-in database of previously computed metagenes whose biological interpretation was already established. The results of this comparison are summarized in interactive html-based tables and JavaScript-based plots, which greatly facilitate exploration of the results. BIODICA also provides some original ways to analyze the metagenes using projection on top of molecular maps (such as InfoSig-Map,(Cantini *et al.*, 2018)). Finally, BIODICA implements the algorithm for constructing RBH graphs, used in the current study.

## 3 Results

Once steps 1 and 2 have been performed as discussed in the Methods section, we obtained the RBH graphs visualized in Figure 2. The nodes of these graphs are the metagenes obtained by the different MFs (A. Stabilized ICA, B. NMF, C. PCA and D. ICA’), different shapes and colors are used to distinguish the different transcriptomic datasets. The topological structure of the obtained graphs is substantially different.

**Fig. 2.**
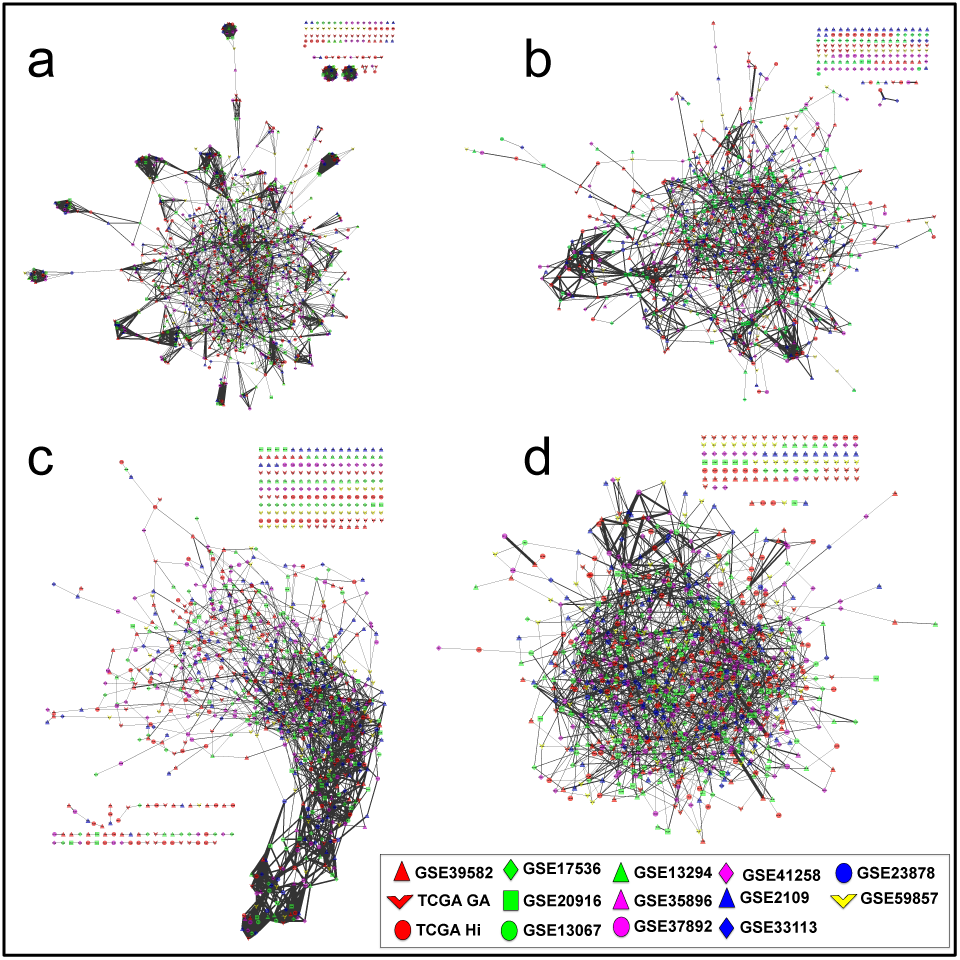
RBH graphs of state-of-art MFs. The RBH graphs obtained in CRC by a) Stabilized ICA; b) NMF; c) PCA and d) ICA’ are here reported.

The RBH graph of Stabilized ICA (Figure 2a) is characterized by clearly pronounced pseudo-cliques and less disconnected nodes in respect to the others. NMF (Figure 2b) has some areas of densely connected nodes but overall this topological aspect is less pronounced in the graph of NMF in respect to the one of Stabilized ICA. The graph of PCA (Figure 2c) reflects the hierarchical structure of the principal components (PC). A densely connected area can be indeed identified in the lower part of the graph, where the first, second and third PCs are localized. This topological organization is lost when going toward higher-order components.

Finally, the graph of ICA’ (Figure 2d) has a surprisingly divergent structure in respect to the one of Stabilized ICA, with a much lower number of pseudo-cliques. This last result suggests that the protocol used to apply ICA has a strong impact in the obtained RBH graph.

The qualitative characteristics discussed above will be extensively tested in the next sections, devoted to the comparison of the measures defined as step 4 of our framework.

### 3.1 Controlling the number of false discoveries

First, we want to assess the ability of the different MFs in producing false positives, or equivalently we want to evaluate how many components identified by the various MFs are prone to noise. Having multiple independent transcriptomic datasets from the same biological condition (in our case CRC), we can expect the same biological factors to be dominating the MF results in at least few datasets. As a consequence, a metagene should find a RBH in at least one other dataset. We thus considered that, if this does not happen, that metagene/node is with high probability a false positive result.

To measure the number of false positive results in the outputs of the different MFs we evaluated the number of disconnected nodes/metagenes. As shown in Figure 3A, Stabilized ICA with 65 disconnected metagenes outperforms other approaches. In particular, both NMF and PCA obtained respectively 129 and 173 disconnected nodes indicating that approximately 13-18% of the signals identified by these MFs are false positives results. Finally, concerning cophNMF, it obtained 12% of disconnected nodes against the 6.7% obtained by Stabilized ICA (see Supplementary Table 2).

**Fig. 3.**
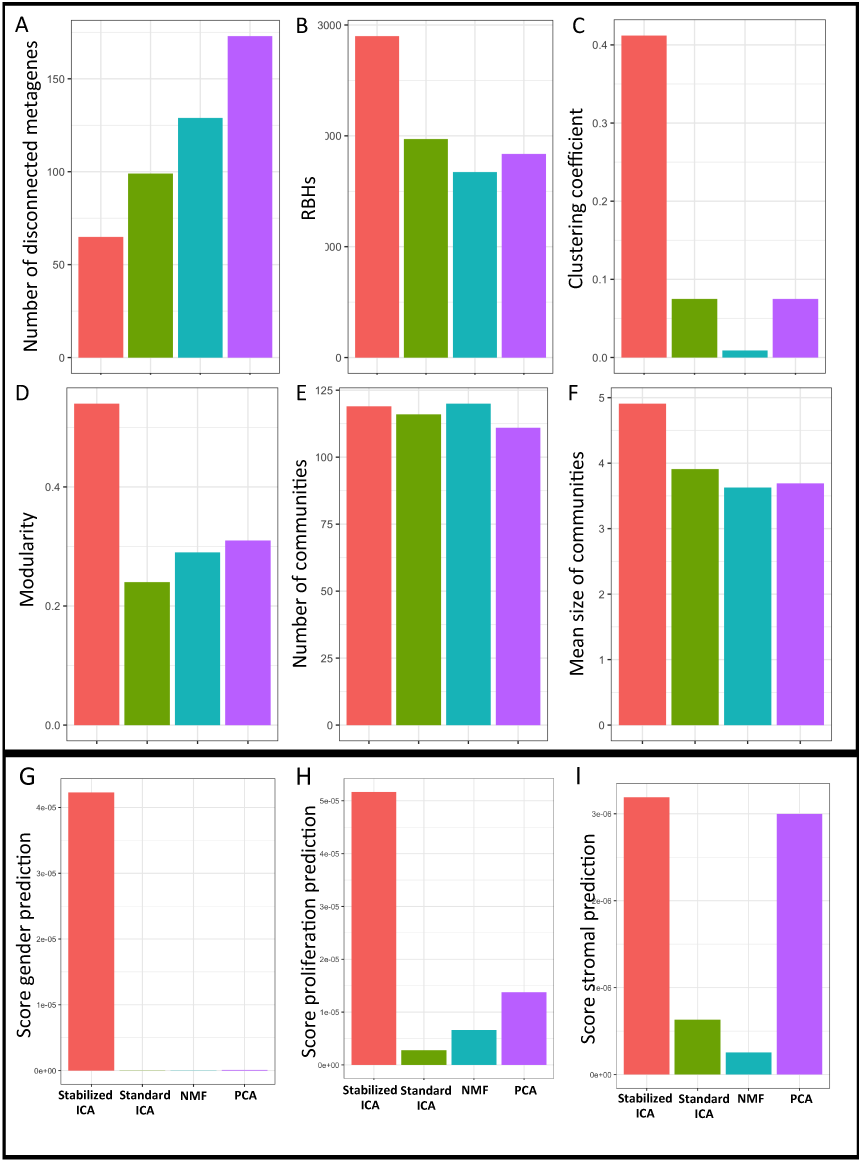
Comparison performances MFs. Different measures are here plotted for the comparison of the various MFs: Stabilized ICA (red), ICA’ (green), NMF (blue) and PCA (violet).

### 3.2 Reproducibility of the identified components

In order to evaluate the reproducibility of the metagenes output of the different MFs we computed the number of links in their RBH graphs. Indeed, working with 14 CRC datasets, in an optimal scenario a metagene should find 13 RBHs corresponding to the metagenes that reflect the same biological factor in all the remaining 13 CRC transcriptomic datasets. In reality, this is not always the case given that a biological factor can be underrepresented in some datasets due to the choice of the samples or to their number. However, higher is the number of RBHs lower is the deviation of the performances of a MF approach in respect to the optimal scenario. As shown in Figure 3B, Stabilized ICA with 2900 RBHs, identifies approximately 1000 more RBHs in respect to the alternative approaches, strikingly outperforming them. Finally, concerning cophNMF, it obtained a 246 RBHs against the 2900 obtained by Stabilized ICA (see Supplementary Table 2).

### 3.3 Presence of a cluster graph-like topology in the RBH graph

Concerning then the topological structure of the RBH graph, the best MF algorithm should derive a cluster-graph like graph, i.e. a disjoint union of cliques. Indeed as discussed above an optimal MF algorithm should find a component for each relevant biological factor underlying the transcriptome. Working with various transcriptomic datasets obtained from the same disease (CRC), those components associated to the same biological factor should cluster together forming a pseudo-clique. The final structure of the optimal RBH graph should be thus composed of various pseudo-cliques sparsely connected one to each other.

In order to verify how the RBH graphs resulting from the different MF approaches are close to this optimal topology, we considered four well-established measures: (i) clustering coefficient; (ii) modularity; (iii) number of communities and (iv) average size of the communities. The first two measures evaluate how evident is the presence of communities in the graph. The average size and number of the communities are instead used to evaluate how consistently each MF algorithm merges components obtained from different datasets. From the results reported in Figure 3C-F the superior performances of Stabilized ICA with respect to alternative approaches can be clearly appreciated. Especially the clustering coefficient and modularity are strikingly higher in Stabilized ICA in respect to ICA’, NMF and PCA. Only for the number of communities NMF obtains a similar, even if slightly lower, value in respect to Stabilized ICA. As shown in Supplementary Table 2, also concerning the topology of the RBH graph the performances of NMF do not improve if considering cophNMF.

### 3.4 Biological content and selectivity of the identified components

Finally, we checked if the communities identified in the RBH graph were effectively associated to specific biological factors. In particular, we tested the ability of the communities of the different MFs in predicting three biological factors that are expected to influence transcriptomic profiles of CRC: patient gender, proliferation status of a tumor and the level of stromal infiltration. For this test we performed a regression analysis of the metasamples obtained from the different MFs.

The gender annotation is composed of discrete values M/F obtained from the available clinical annotations: in this case, we thus performed a logistic regression. Proliferation was evaluated averaging the expression of the genes belonging to a well-known proliferation signature (Giotti *et al.*, 2017) and it is thus a vector of continuous weights. Finally, stromal infiltration was estimated using the average expression of the genes belonging to the stromal signature of ESTIMATE tool (Yoshihara *et al.*, 2013).

The results of this first test are summarized in Figure 3 G-I. We focused on the community that predicted the best the specified biological signal. The community was selected as the one with the highest percentage P of metasamples whose regression on the biological signal was significant. We used three parameters commonly used to evaluate the quality of a linear regression: R^2^, Bayesian information criterion (BIC) and Akaike’s information criterion (AIC). We finally define a score to combine them in a single value as (P*R2) / (BIC*AIC). The higher this score the stronger is the association between the community and the biological factor. Indeed a good regression would correspond to R^2^ value near to 1 and low BIC and AIC values. Such scores are reported in Figure 3 G-I. The specific values obtained by the single scores are reported in Supplementary Table 3. As shown in Figure 3 G-I and Supplementary Table 2, Stabilized ICA is the approach that better approximates all three tested biological factors. In particular, NMF does not identify any component that can significantly predict the gender signal.

We then investigated the selectivity of such predictions, meaning the ability of the MF approach to define a clear one-to-one association between a biological signal and a component. To test for the selectivity of the different MFs we focused on the components obtained on the GSE39582 dataset (see Supplementary Table 1) and considered the R^2^ obtained in the previously computed regressions by all the 100 components. As shown in Supplementary Figure 3, Stabilized ICA resulted to be far more selective than the alternative MFs. In particular for all the three biological factors (gender, proliferation and stromal infiltration) Stabilized ICA found only one component strongly associated to them. On the opposite, NMF and ICA’ identified multiple components with similar regression performances. Finally PCA resulted to be selective in stromal infiltration and proliferation prediction. However, PC1 was the component predicting simultaneously both signals, confirming the already observed limitation of PCA of conflating multiple biological processes into a single component.

### 3.5 Impact of the technical platform on the results obtained by the different MFs

To finally test the impact of the transcriptome profiling platforms on the results of the various MF algorithms we took advantage of four ovarian cancer transcriptomic datasets available from TCGA (Bell *et al.*, 2011). These four datasets have been profiled with four different platforms: Affymetrix U133, Agilent and Affymetrix HuEx, plus RNAseq. 418 samples are in common among all four datasets. This resource is optimal to evaluate if the profiling platform affects the results of the various MFs. Indeed having four datasets composed of the same samples we are sure that no biological variability is present across them. In the optimal scenario, all the metagenes of an MF algorithm should find a RBH with a metagene of the other three datasets. To thus assess which MF algorithm is less prone to technical noise, we applied to each MF step 1 and 2 of our new framework. We thus decomposed the four ovarian cancer datasets in 100 components and we then reconstructed the RBH graph. We finally checked the number of RBH links of the different MFs and their average absolute correlation. Stabilized ICA resulted to perform better than alternative approaches also in this case, with 390 links and average correlation of 0.396 (see Supplementary Table 4 for all the results).

### 3.6 Stabilized ICA identifies consistent biological insights on CRC in respect to previous knowledge

In previous sections we showed that Stabilized ICA performs better than alternative approaches according to multiple measures of practical interest for highthroughput data analysis. We thus now concentrate more deeply on the biological insights that can be derived from the RBH graph of this MF algorithm. To this aim we added to the analysis other four datasets: single-cell RNAseq from normal and tumoral CRC tissue (Li *et al.*, 2017), Patient-derived Xeno graft (PDX) CRC Models and liver metastasis (LM) (Isella *et al.*, 2017). We then biologically annotated the communities of the RBH graph by using meta-metagenes and metasamples according to the procedure described in the Methods section. The meta-metagenes obtained for the communities of Stabilized ICA are reported in Supplementary Table 5 and represent a useful resource for further analyses.

Figure 4 and Supplementary Table 5 report the RBH graph of Stabilized ICA and the main biological informations extracted from it. Four main categories of biological factors can be distinguished in the graph: factors intrinsic to the tumor, microenvironment signals, technical signals, effects of small groups of genes and finally unknown factors. Concerning the tumor-specific factors, some communities were found to be associated to core tumoral functions, such as proliferation, inflammation, stemness, interferon response and mitochondria. Other tumorspecific communities resulted instead to be associated to CRC-specific tumoral signals, such as MSI/MSS (microsatellite instability/microsatellite stable), goblet cells (a differentiated cell of the colon) and KRAS mutation. Finally, one community was found to be related to chromatin silencing and histones. The stromal communities instead include microenvironment signals, such as cancer-associated fibroblasts (CAFs), smooth muscle, immune, complement system and B-cells. Of particular interest is the identification of a community related to B-cells whose association to this cell type was evident not only using the MSigDB signatures (as discussed in Methods), but also from single-cell data. Looking indeed at the metasample of the component obtained from single-cell data and associated to this community we could clearly see B-cells being the most contributing in this component. The technical factors included instead GC-content and gender. Finally, 10 communities have been found to be associated with small groups of genes. In this last case, the meta-metagenes associated to these communities contained few genes having a much higher weight than the others.

**Fig. 4.**
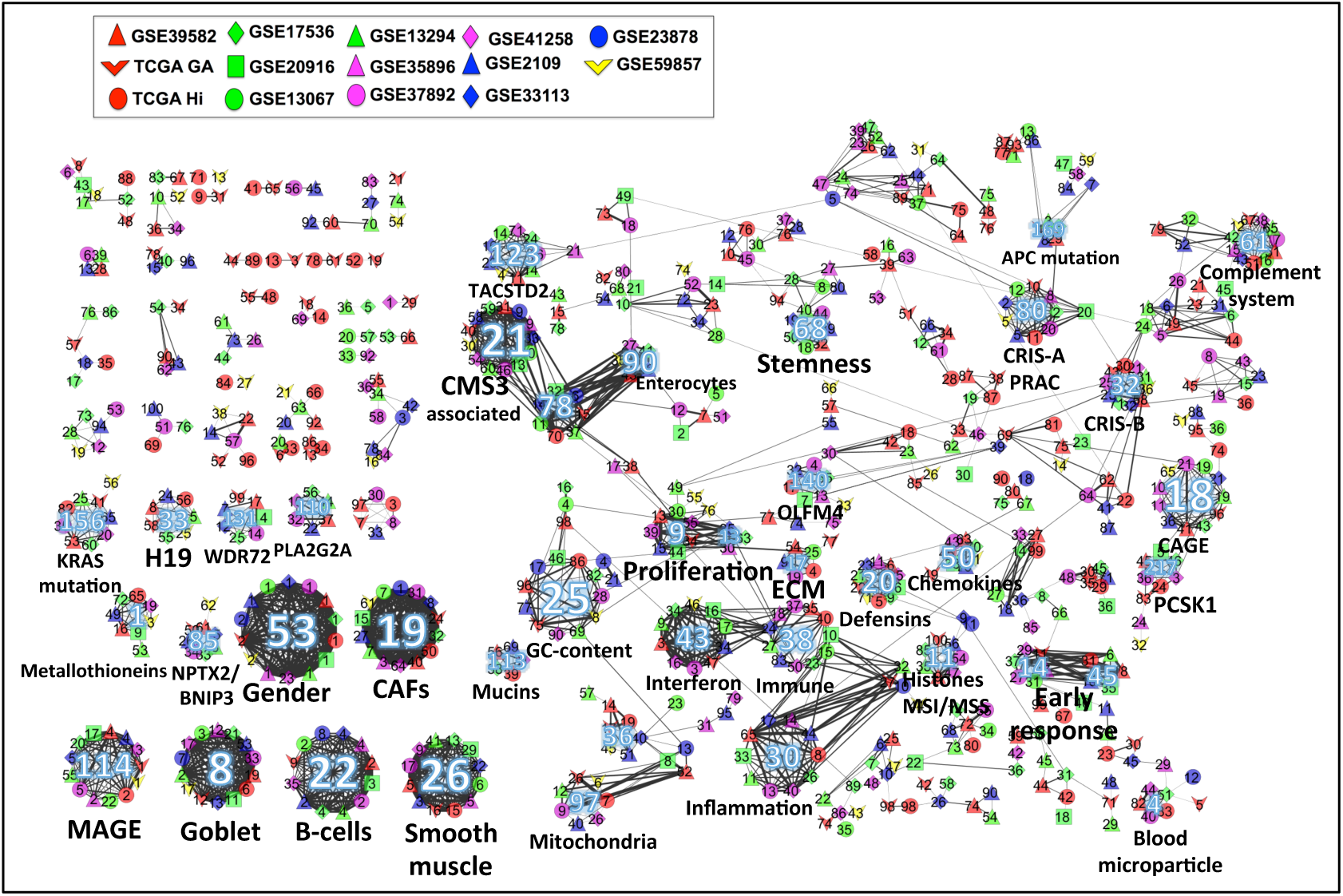
RBH graph of Stabilized ICA with the main biological annotations. The node colors indicate the dataset from which the components have been computed. The edge thickness indicates the magnitude of the correlation. Communities with more than six elements are marked with an integer number. For details on the community annotations see Supplementary Table 5.

Concerning the association with the predefined CRC Consensus Molecular Subtypes (CMS) we could clearly match CMS1 with our immune component, concordantly to what previously observed. Communities associated to CMS3 and CMS4 were also identified. Of note, the CMS4 subtype resulted from our analysis to be associated to both smooth muscles and CAFs. A strong CAFs infiltration had been already observed in this CRC subtype (Isella *et al.*, 2015; Guinney *et al.*, 2015).

## 4 Discussion

In this manuscript we compared the three most commonly used matrix factorization methods for their ability to detect reproducible and biologically interpretable signals in independent transcriptomic datasets of the same cancer type (CRC). For one of the methods, Independent Component Analysis, we also compared two protocols of its application to transcriptomic data, named ICA’ and Stabilised ICA. We designed a new framework based on the concept of Reciprocal Best Hit (RBH), borrowed from evolutionary bioinformatics. Applying our new framework to state-of-art MFs, we can definitively conclude that Stabilised ICA is able to deliver more reproducible and interpretable results compared to other methods on transcriptomic datasets.

This advantageous features of Stabilised ICA can be explained by the fact that the statistical model lying behind ICA matches better the geometrical structure of transcriptomic datasets in the multi-dimensional gene expression space, compared to other methods. Minimizing independence of biological mechanisms represented by metagenes rather than independence of biological samples, results in better metagene interpretability and matching between datasets. Using multiple runs of ICA for stabilisation and prioritizing stable components also significantly contributes to improving their reproducibility (see (Kairov *et al.*, 2017). By contrast, PCA components appear to systematically mix multiple sources of transcriptome variability, reducing interpretability. Also, the higherorder PCA components are regularly not reproducible. The main problem of the NMF metagenes is that in application to gene expression data, most of them are strongly correlated with average gene expression. As a result, even if NMF is able to delineate an important region of high level and variability in the gene expression space, the individual NMF components are rarely specifically and selectively associated with biological factors, which ruins both their reproducibility and interpretation.

We demonstrated that the meta-analysis of the results of Stabilised ICA, based on constructing the RBH graph, provides a biologically rich image of the signals shaping tumoral transcriptomes and their interconnection. Pseudo-cliques, existing in the RBH graph, whose meaning can be compared to the Clusters of Orthologous Genes (COGs) in evolutionary bioinformatics, can be matched to previously known and/or expected highly reproducible biological signals (such as proliferation and immune infiltration) but also highlights novel biological mechanisms which require further investigation and interpretation. The metagenes obtained through application of MF methods can be compared to other methods, sharing similar spirit. In particular, attractor metagenes were suggested in order to serve as surrogates of cancer phenotypes (Cheng *et al.*, 2013). Attractor metagenes were used as variables in the DREAM Challenge winning approach for predicting breast cancer clinical outcome (Margolin *et al.*, 2013). We find ICA-based framework for identifying metagenes more computationally elegant and potentially producing less false positive signatures; however, further study is required to compare the results of both approaches and their computational performances. INSPIRE method uses the latent variable approach to infer modules of co-expressed genes and the dependencies among the modules from multiple expression datasets that may contain different sets of genes (Celik *et al.*, 2016). Therefore, INSPIRE shares general objectives of MF-based meta-analysis but significantly differs in terms of methodology. For example, INSPIRE is based on the assumption of Gaussianity in the data distributions and uses disjoint module definitions rather than metagenes, where each gene can contribute to several biological functions.

Lastly, here we compared MF methods in application to cancer transcriptomic datasets. However, the suggested approach can be easily extrapolated to other data types (methylomic, proteomic) or other fields of research collecting massive transcriptomic datasets (such as drug screenings).

## Acknowledgements

We thank Urszula Czerwinska and Alessandro Greco for testing the BIODICA software. We thank Askhat Molkenov for developing the BIODICA GUI. We thank Justin Guinney and the whole Colorectal Cancer Subtyping Consortium (CRCSC) for supporting the idea and providing the normalized datasets.

---

**Supplementary Figure 1.**
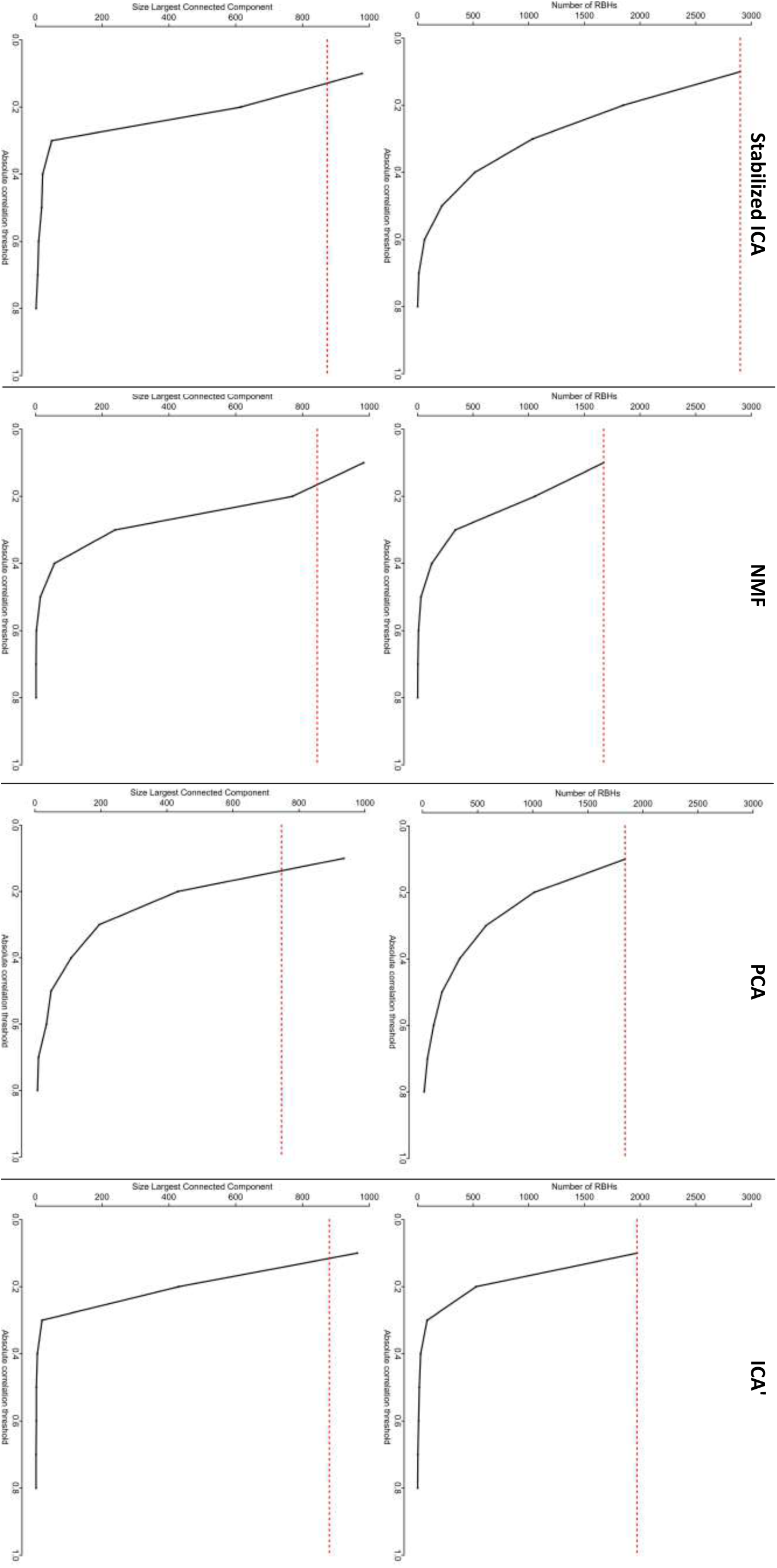
Comparison between RBH graph (red) and correlation graph (black) using different correlation thresholds. For each of the four MFs the behavior of the number of RBHs (top) in the network and the dimension of the largest connected component (bottom) are reported for different values of the correlation threshold.

**Supplementary Figure 2.**
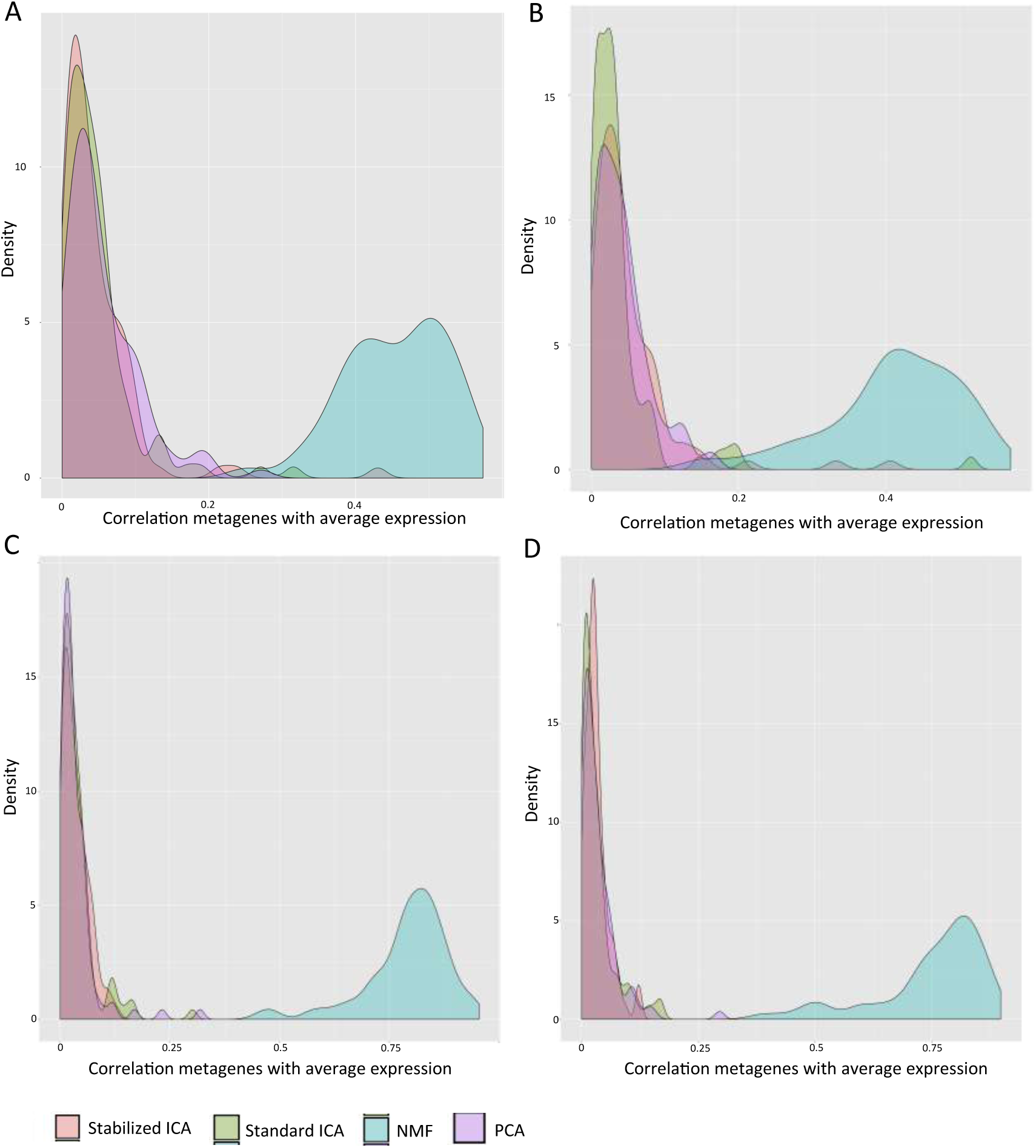
Distribution of the correlations of the MF metagenes in the four bigger transcriptional datasets A) GSE39582 B) GSE2109 C) TCGA-GA D) TCGA Hi-seq. Different colors denote the different MD methods: red for Stabilized ICA, green for ICA’, blue for NMF and violet for PCA.

**Supplementary Figure 3.**
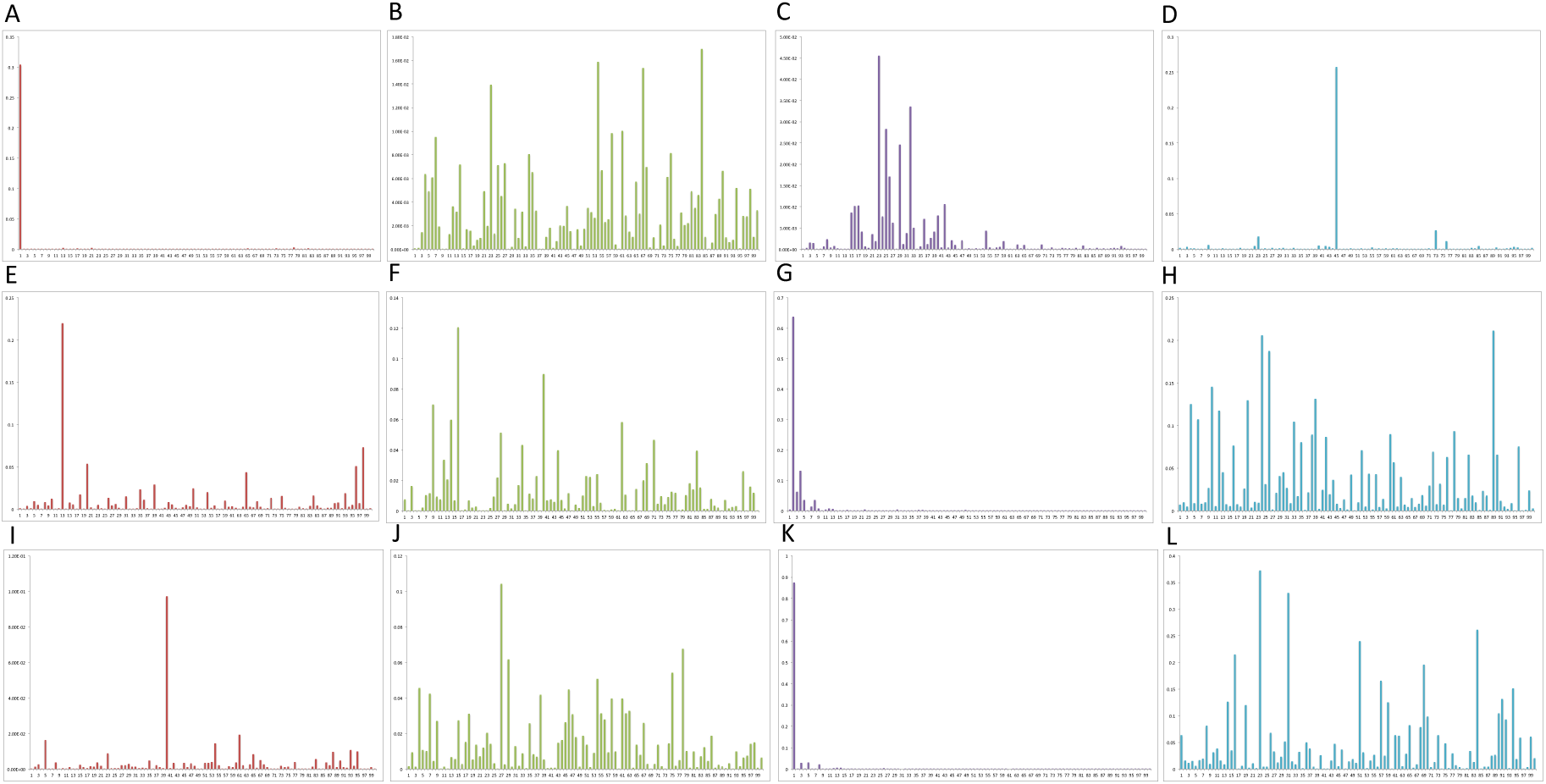
R^2^ values of the regression of the 100 metagenes computed in GSE39582 on the gender (A-D), proliferation (E-H) and stromal infiltration (I-L) signals. Different colors denote the various MF algorithms used to compute the 100 metagenes: red for Stabilized ICA, green for ICA’, violet for PCA and blue for NMF. The x-axis reports the metagenes in ascending order, while y-axis reports the R^2^ value.

**Supplementary Table 1.** List of CRC datasets considered in this comparison. For each dataset, the GEO identifier, the number of samples and the number of computed components is here reported

|  |  |  |
| --- | --- | --- |
| **Dataset code** | **number of samples** | **number of components** |
| GSE23878 | 35 | 18 |
| GSE35896 | 62 | 31 |
| GSE13067 | 74 | 37 |
| GSE33113 | 90 | 45 |
| GSE20916 | 100 | 50 |
| GSE37892 | 130 | 65 |
| GSE13294 | 155 | 78 |
| GSE59857 | 155 | 78 |
| GSE17536 | 177 | 89 |
| GSE41258 | 187 | 94 |
| TCGA GA | 264 | 100 |
| GSE2109 | 277 | 100 |
| TCGA HI | 332 | 100 |
| GSE39582 | 566 | 100 |

**Supplementary Table 2.**
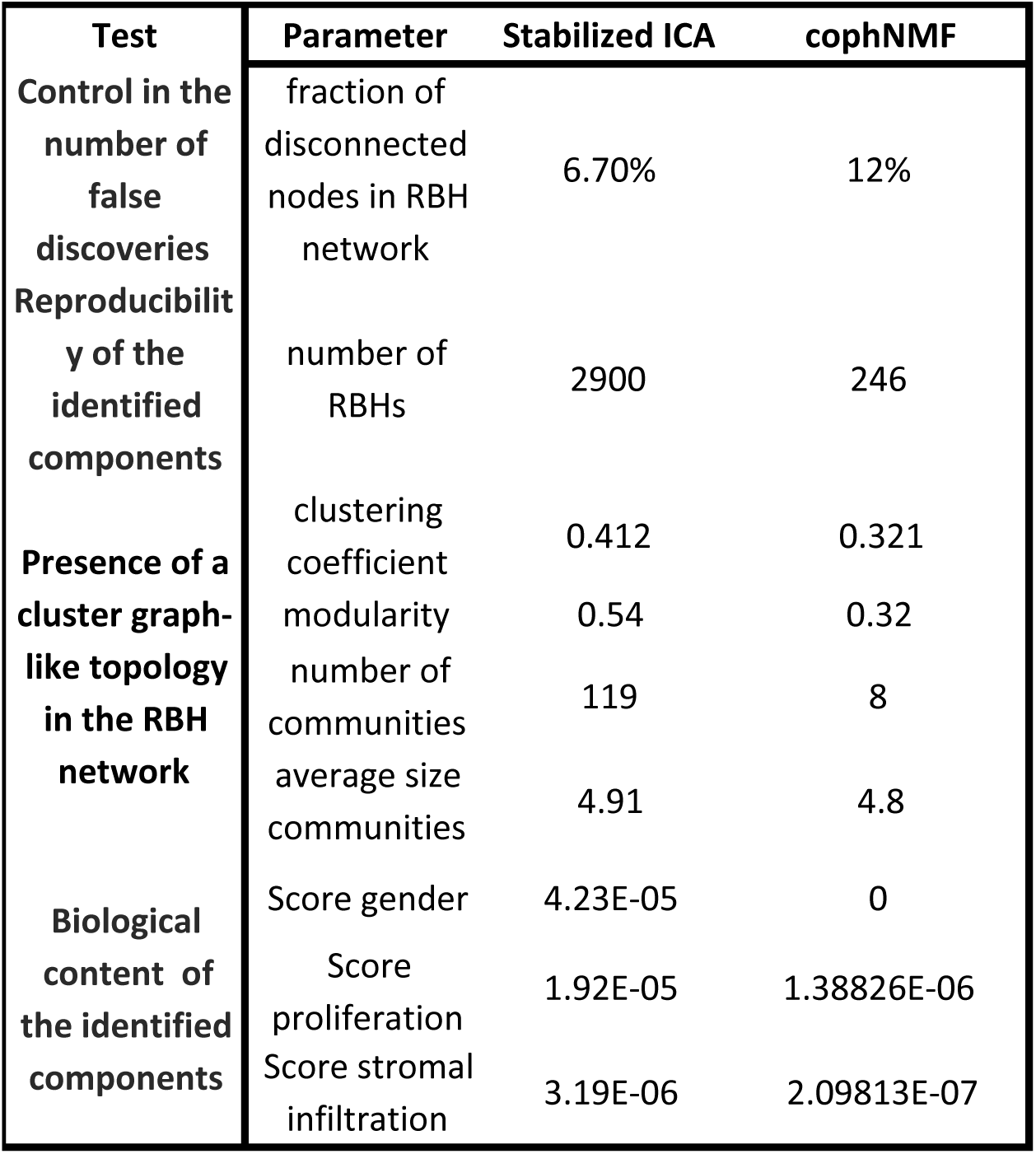
Comparison between the performances of Stabilized ICA and cophNMF. The Results of Stabilized ICA and cophNMF in the different tests reported in the results are here summarizedIn the first columns the names of the main results sections are reported.

**Supplementary Table 3.**
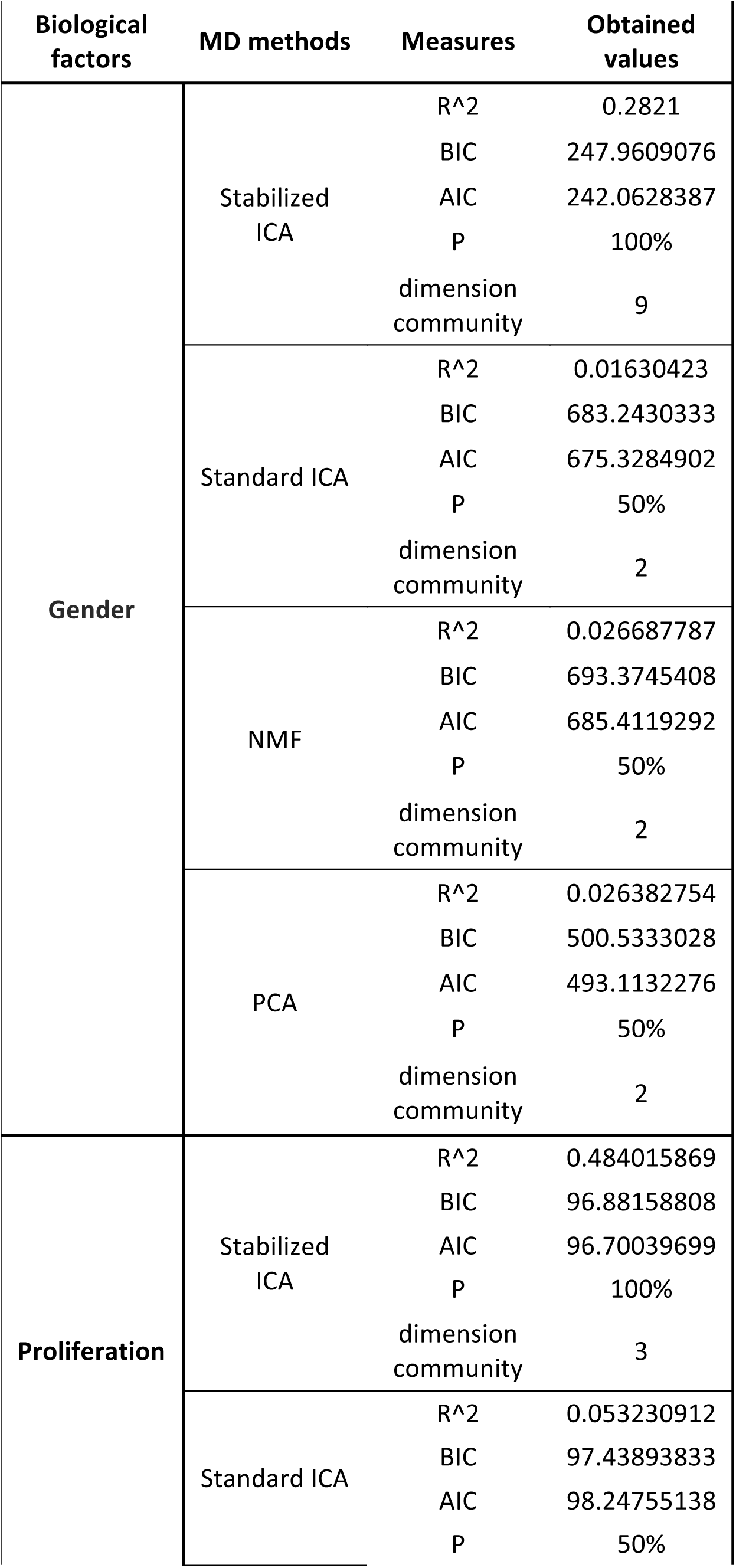
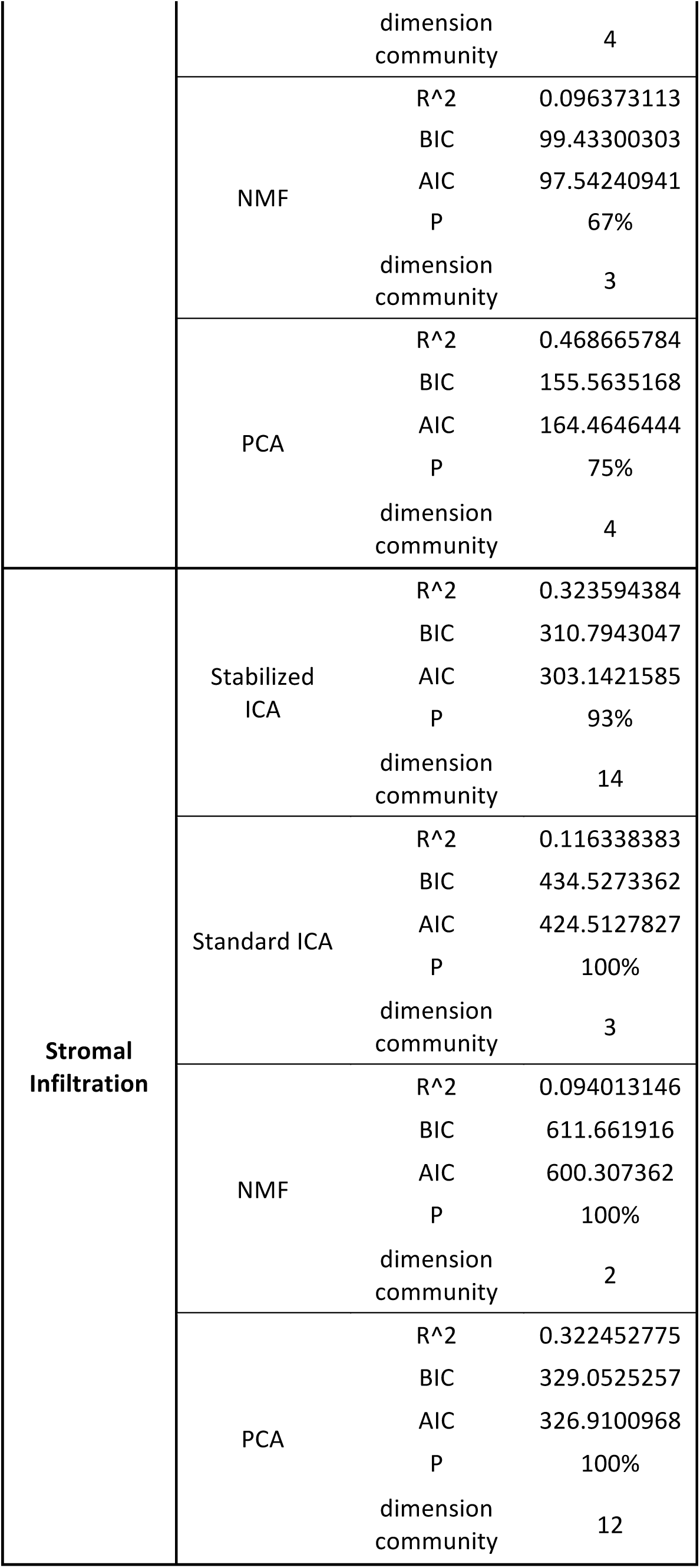
Comparison performances in the regression of the three biological signals: gender, proliferation and stromal infiltration by Stabilized ICA, ICA’, NMF and PCA. The table reports the values summarized in Figure 4: R^2, BIC, AIC, P percentage of components in the community with a significant regression and dimension of the corresponding community.

**Supplementary Table 4.**
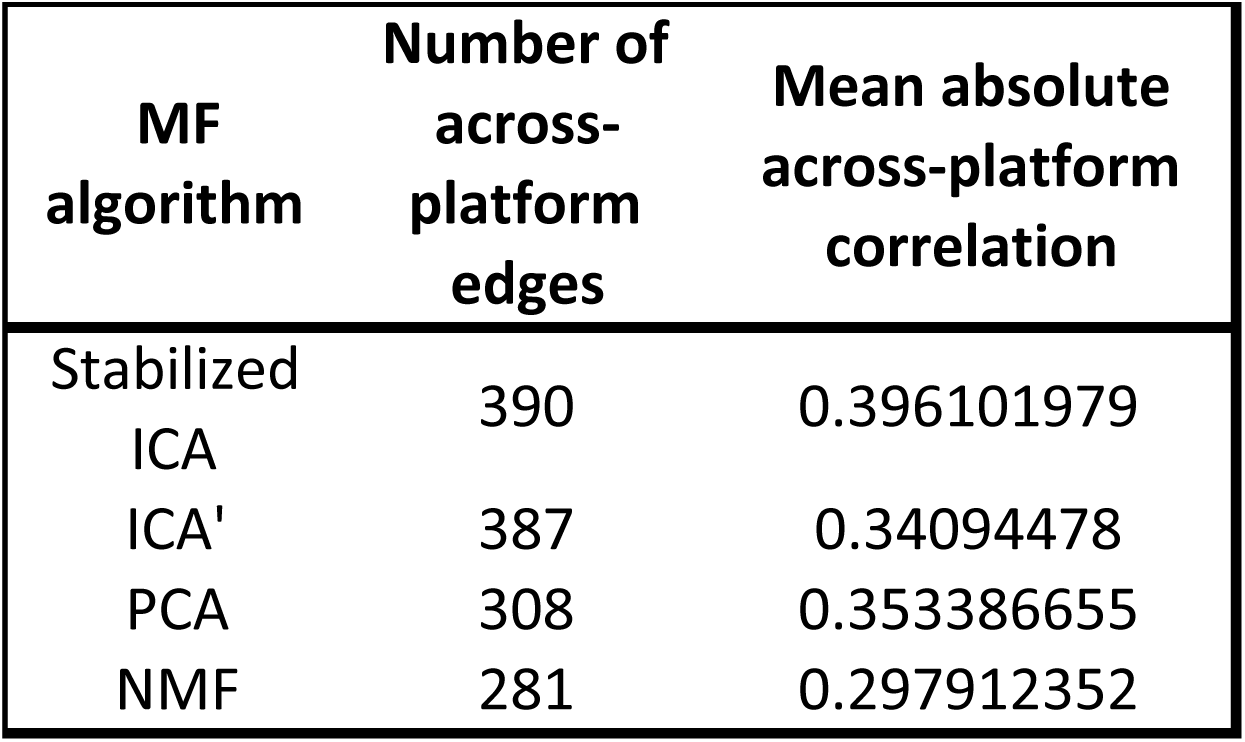
Comparison of acrossplatform reproducibility of the components obtained by the different MF algorithms. For each MF reported in the first column, the number of RBHs connecting metagenes from different platforms (second column) and the average correlation of such links (third column) is reported.

**Supplementary Table 5.**
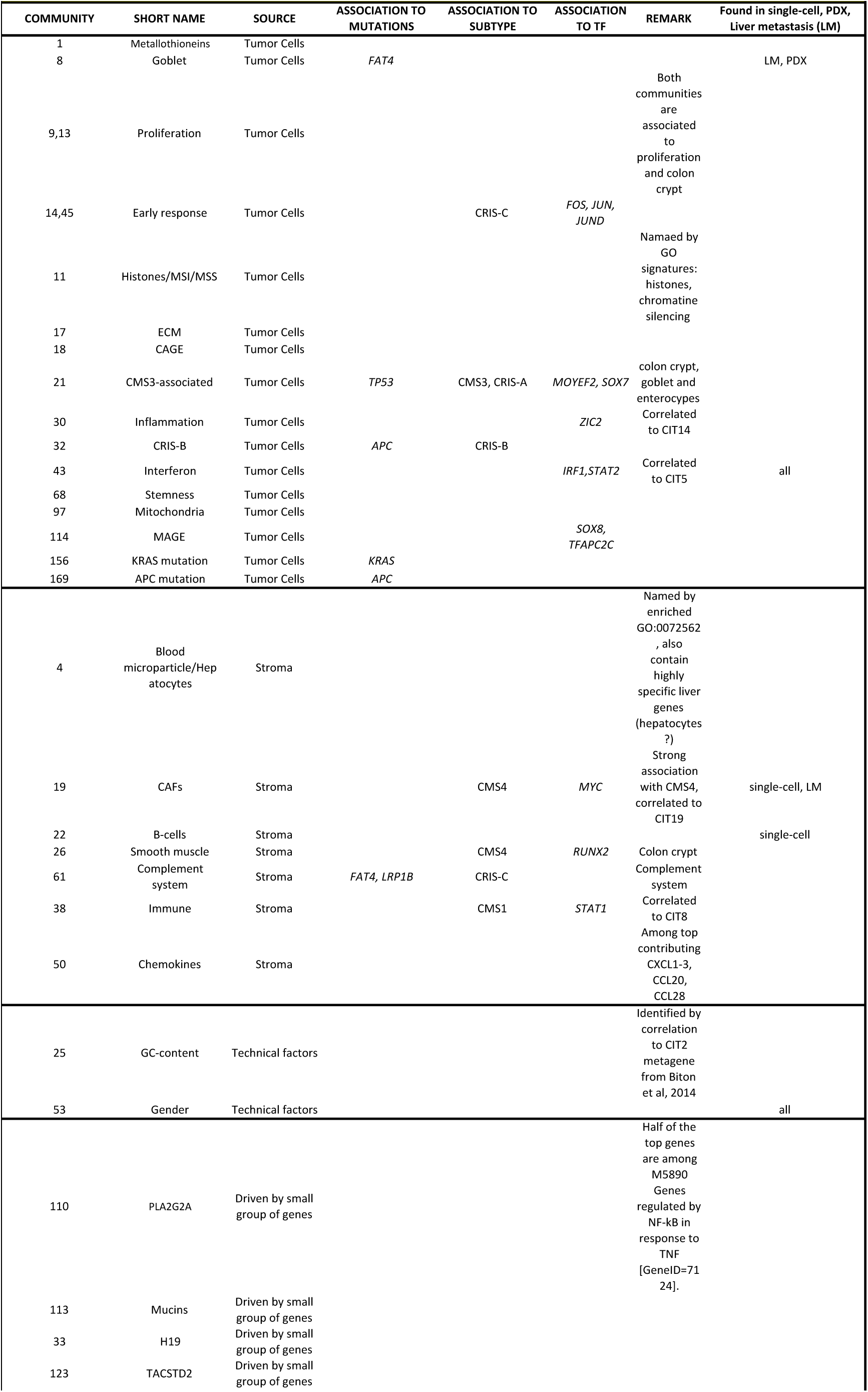
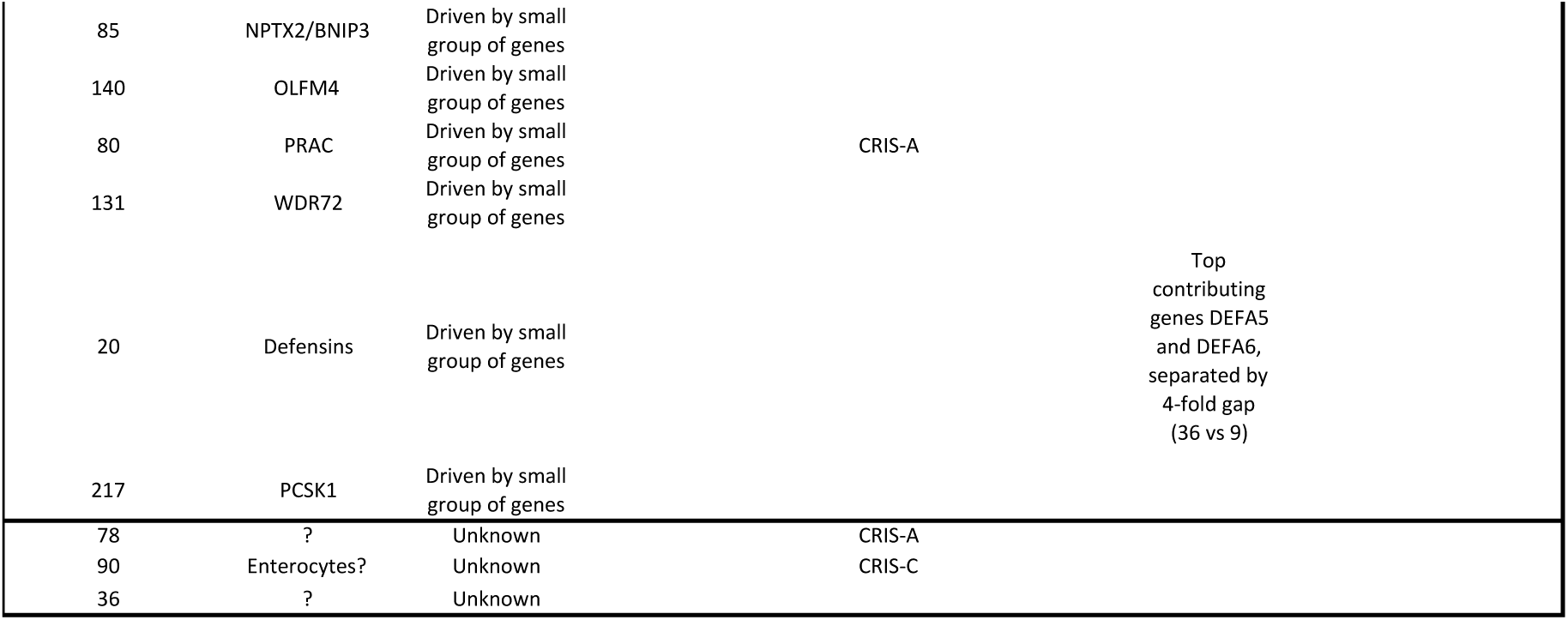
The table reports the main biological annotations available for the communities of Figure 5. Communities have been obtained using MCL on the Stabilized ICA PBH network, these communities are numbered with integer values and their number is reported in the first column. The second column reports the short name of each community, corresponding to the biological process most significant in the community. In the third column, the four main sources of the identified biological factors are reported. Several biological factors strongly associated with a community corresponded to mutations found in colorectal cancer (column 4). Several communities were strongly associated with a subtype of colorectal cancer described in (Guinney et al. 2015) (CMS1, CMS3, CMS4) or in (Isella et al. 2016) (CRIS-A, CRISB, CRIS-C, CRIS-D, CRIS-E) (column 5). For several communities, the most contributing genes correspond to target genes of a transcription factor (column 6). Several communities have been found in other cancers and described in (Biton et al. 2014). In this paper, the communities were named CX. Finally, in column 8 the presence of the community also in single-cell, PDX and liver metastasis is annotated.

